# Intracellular NASPM allows an unambiguous functional measure of GluA2-lacking calcium-permeable AMPA receptor prevalence

**DOI:** 10.1101/2021.02.18.431828

**Authors:** Ian D. Coombs, Cécile Bats, Craig A. Sexton, Stuart G. Cull-Candy, Mark Farrant

## Abstract

Calcium-permeable AMPA-type glutamate receptors (CP-AMPARs) contribute to many forms of synaptic plasticity and pathology. They can be distinguished from GluA2-containing calcium-impermeable AMPARs by the inward rectification of their currents, which reflects voltage-dependent block by intracellular spermine. However, the efficacy of this weakly permeant blocker is differentially altered by the presence of AMPAR auxiliary subunits – including transmembrane AMPAR regulatory proteins, cornichons and GSG1L – that are widely expressed in neurons and glia. This complicates the interpretation of rectification as a measure of CP-AMPAR expression. Here we show that inclusion of the spider toxin analogue 1-naphthylacetyl spermine (NASPM) in the intracellular recording solution results in complete block of GluA1-mediated outward currents irrespective of the type of associated auxiliary subunit. In neurons from GluA2-knockout mice expressing only CP-AMPARs, intracellular NASPM, unlike spermine, blocks all outward synaptic current. Thus, our results identify an unambiguous functional measure, sensitive solely to changes in CP-AMPAR prevalence.

## Introduction

AMPA-type glutamate receptors (AMPARs) mediate the fast component of excitatory postsynaptic currents (EPSCs) throughout the mammalian brain (Baranovic and Plested, 2016; Greger et al., 2017; Traynelis et al., 2010) and their regulation allows lasting changes in synaptic strength that are essential for normal brain function (Huganir and Nicoll, 2013). AMPARs are nonselective cation channels that exist as homo- or heterotetrameric assemblies of the homologous pore-forming subunits GluA1−4, encoded by the genes *Gria1-4*. While a majority of central AMPARs are GluA2-containing di- or tri-heteromeric assemblies (Lu et al., 2009; Wenthold et al., 1996; Zhao et al., 2019), those lacking GluA2 constitute an important functionally distinct subtype. Editing of *Gria2* pre-mRNA at codon 607 results in the substitution of a genetically encoded glutamine (Q) with an arginine (R). This switch from a neutral to a positively charged residue in the pore-forming loop causes receptors containing Q/R edited GluA2 to have a greatly reduced Ca^2+^ permeability compared to those lacking GluA2 (Burnashev et al., 1992; Kuner et al., 2001; Sommer et al., 1991). As this mRNA editing is essentially complete, AMPARs are commonly divided into GluA2-containing Ca^2+^-impermeable (CI-) and GluA2-lacking Ca^2+^-permeable (CP-) forms (Bowie, 2012; Burnashev et al., 1992; Cull-Candy et al., 2006).

Aside from allowing Ca^2+^ flux, CP-AMPARs differ from CI-AMPARs in having greater single-channel conductance (Benke and Traynelis, 2019; Swanson et al., 1997) and susceptibility to voltage-dependent block by the endogenous intracellular polyamines spermine and spermidine (Bowie and Mayer, 1995; Donevan and Rogawski, 1995; Kamboj et al., 1995; Koh et al., 1995). They can also be blocked in a use-dependent manner by extracellular application of the same polyamines (Washburn and Dingledine, 1996) or various exogenous organic cations, such as the spermine analogue *N*-(4-hydroxyphenylpropanoyl)-spermine (HPP-SP; Washburn and Dingledine, 1996), the polyamine-amide wasp toxin philanthotoxin-4,3,3 (PhTx-433; Washburn and Dingledine, 1996), the spider toxin analogue 1-naphthylacetyl spermine (NASPM; Tsubokawa et al., 1995), and the dicationic adamantane derivative IEM-1460 (Magazanik et al., 1997).

Block of CP-AMPARs by intracellular polyamines results in inwardly- or bi-rectifying current-voltage relationships. This characteristic rectification – seen during whole-cell patch-clamp recordings in the presence of residual endogenous polyamines or added exogenous spermine – has been utilised extensively to identify the presence of CP-AMPARs in neurons. Importantly, this approach has enabled the identification of cell- and synapse-specific CP-AMPAR expression (Koh et al., 1995; Toth and McBain, 1998), changes in CP-AMPAR prevalence during development (Brill and Huguenard, 2008; Kumar et al., 2002; Soto et al., 2007; Wang and Gao, 2010), and roles for CP-AMPARs in multiple synaptic plasticities, including long-term potentiation (LTP) and depression (LTD) (Fortin et al., 2010; Lamsa et al., 2007; Liu and Cull-Candy, 2000; Mahanty and Sah, 1998; Plant et al., 2006; Sanderson et al., 2016). In addition, its use has revealed CP-AMPAR changes associated with ischaemia (Dixon et al., 2009; Liu et al., 2004; Peng et al., 2006), brain trauma (Korgaonkar et al., 2020) and glaucoma (Sladek and Nawy, 2020), and in circuit remodelling associated with chronic disorders, including fear-related behaviours (Clem and Huganir, 2010; Liu et al., 2010), drug addiction (Bellone and Luscher, 2006; Conrad et al., 2008; Lee et al., 2013; Scheyer et al., 2014; Van den Oever et al., 2008), and neuropathic or inflammatory pain (Goffer et al., 2013; Katano et al., 2008; Park et al., 2009; Sullivan et al., 2017; Vikman et al., 2008).

Native AMPARs co-assemble with various transmembrane auxiliary subunits that influence receptor biogenesis, synaptic targeting and function (Greger et al., 2017; Jackson and Nicoll, 2011a; Schwenk et al., 2019). Importantly, several of these, including transmembrane AMPAR regulatory proteins (TARPs), cornichons, and germ cell-specific gene 1-like protein (GSG1L), have been shown to modify the block of CP-AMPARs by intracellular spermine. In the case of TARPs and CNIHs inward rectification is reduced (Brown et al., 2018; Cho et al., 2007; Coombs et al., 2012; Soto et al., 2007; Soto et al., 2009), whereas with GSG1L the rectification is increased (McGee et al., 2015). This complicates the interpretation of measures of spermine-dependent rectification. Moreover, any change in rectification that might be attributed to changes in the prevalence of CP-AMPARs could reflect changes in auxiliary subunit content.

Here we show that intracellular NASPM, PhTx-433 and PhTx-74 specifically and voltage-dependently block CP-AMPARs. Unlike spermine, which is permeant, these blockers allow minimal outward current at positive potentials. At +60 mV, 10 μM intracellular NASPM produces a use-dependent block but at 100 μM it fully blocks outward currents in a use-independent manner. Importantly, this block is unaffected by auxiliary subunit content. Accordingly, in TARP γ2- and γ7-expressing cerebellar stellate cells (Fukaya et al., 2005; Yamazaki et al., 2010) cultured from GluA2-lacking (GluA2^-/-^) mice (Jia et al., 1996), 100 μM intracellular NASPM causes full rectification of CP-AMPAR-mediated miniature EPSCs (mEPSCs), something that cannot be achieved with spermine. Together, our results reveal intracellular NASPM to be an ideal tool with which to determine the contribution of CP-AMPARs to AMPAR-mediated currents.

## Results

### Intracellular polyamine toxins block CP-AMPARs

To determine whether intracellularly applied NASPM or polyamine toxins might offer advantages over spermine for the identification of native CP-AMPARs we examined three compounds: NASPM, which contains the same polyamine tail as spermine, PhTx-433, which has a different distribution of amines, and PhTx-74, which lacks one amine group (**Fig. 1a**). Initially, we recorded currents in outside-out patches from HEK cells transiently transfected with GluA1 alone or with GluA1 and GluA2, to produce homomeric CP- and heteromeric CI-AMPARs, respectively. The receptors were activated by glutamate (300 μM) in the presence of cyclothiazide (50 μM) to minimize AMPAR desensitization and we applied voltage ramps (100 mV/s) to generate current-voltage (*I-V*) relationships. As expected (McGee et al., 2015; Soto et al., 2007), in the absence of intracellular polyamines homomeric GluA1 receptors generated outward currents at positive potentials, showing clear outward rectification, while in the presence of 100 μM spermine, they displayed doubly rectifying responses (**Fig. 1b**). By contrast, when the intracellular solution contained 100 μM NASPM, GluA1 receptors displayed inwardly rectifying responses with negligible current passed at potentials more positive than +20 mV (**Fig. 1b**). Unlike responses from GluA1 alone, currents from GluA1/2 receptors in the presence of NASPM were non-rectifying (**Fig. 1b**). *I-V* plots showed that, when added to the intracellular solution at 100 μM, each blocker conferred marked inward rectification on the currents mediated by GluA1 (rectification index, RI_+60/-60_ 0.02-0.26), but not on those mediated by GluA1/2 (RI_+60/-60_ 0.84-1.30) (**Fig. 1c**). Thus, although differing in structure, they all produced selective voltage-dependent block of the GluA2-lacking CP-AMPARs. Of note, the block by intracellular PhTx-74 was restricted to CP-AMPARs, despite the fact that it produces low affinity block of CI-AMPARs when applied extracellularly (Jackson et al., 2011; Nilsen and England, 2007).

**Figure 1.**
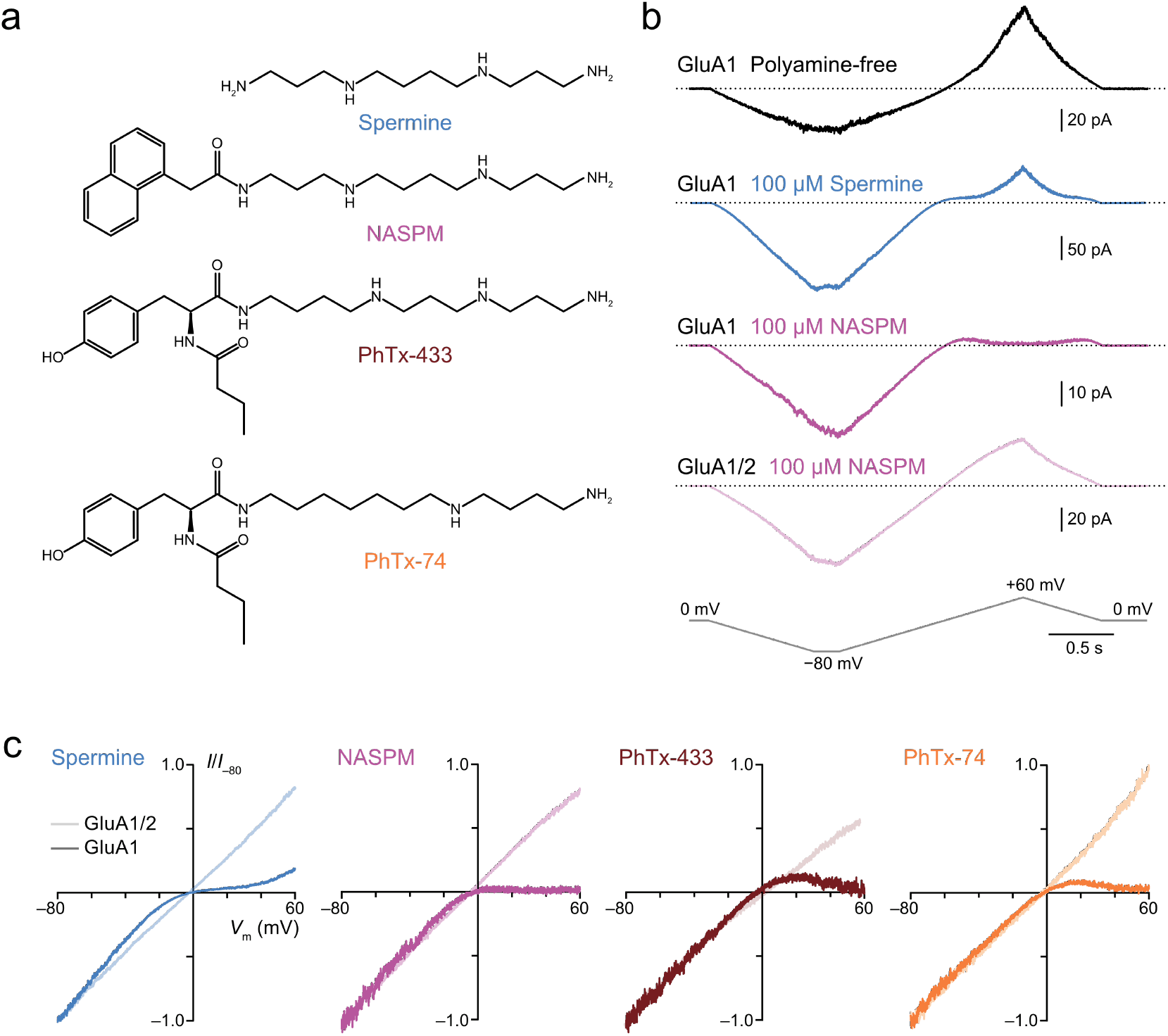
Intracellular NASPM and polyamine toxins specifically block GluA2-lacking CP-AMPARs in a voltage-dependent manner. **a**) The blockers used in this study. **b**) Representative responses activated by 300 μM glutamate and 50 μM cyclothiazide from outside-out patches excised from HEK293 cells expressing either GluA1 or GluA1/2. The voltage was ramped linearly from −80 to +60 mV (100 mV/s). GluA1 displayed outward rectification with a polyamine-free pipette solution. In the presence of 100 μM spermine GluA1 displayed a doubly rectifying relationship, and with 100 μM NASPM GluA1 displayed full inward rectification. GluA1/2 did not rectify in the presence of NASPM. **c**) Normalized and pooled *I-V* relationships for GluA1 and GluA1/2 in the presence of 100 μM intracellular polyamines (*n* = 3-8). For each blocker the RI_+60/-60_ value with GluA1 was less than that with GluA1/2. For spermine, the values were 0.26 ± 0.11 and 1.12 ± 0.07, and the unpaired mean difference was 0.86 [95%CI 0.62, 1.09] (*n* = 5 GluA1 patches and 4 GluA1/2 patches). For NASPM, the values were 0.03 ± 0.02 and 1.10 ± 0.21, 1.07 [0.79, 1.6] (*n* = 4 and 8). For PhTx-433, the values were 0.02 ± 0.04 and 0.84 ± 0.05, 1.25 [0.97, 1.43] (*n* = 4 and 3). For PhTx-74, the values were 0.06 ± 0.09 and 1.30 ± 0.13, 0.82 [0.60, 0.95 (*n* = 4 and 4). **Figure 1—source data** RI_+60/-60_ values from GluA1 and GluA1/γ2 ramp *I-V* relationships (300 μM glutamate) recorded with intracellular spermine, NASPM, PhTx-433 or PhTx-74 (each 100 μM). Fig_1_RI_source_data.csv

We next investigated the concentration- and auxiliary subunit-dependence of the block. Specifically, we generated conductance-voltage (*G-V*) relationships and fit those from inwardly rectifying responses with a single Boltzmann function and those from doubly rectifying responses with a double Boltzmann function (Panchenko et al., 1999). This revealed that as the concentration of added blocker was increased (from 0.1 or 1 μM to 500 μM) there was a progressive negative shift in *V*_b_ (the potential at which 50% block occurs) (**Fig 2a, b**). Plotting *V*_b_ against polyamine concentration (**Fig. 2b**) allowed us to determine the *IC*_50, 0 mV_ (the concentration expected to result in half maximal block at 0 mV) and thus estimate the potency of each polyamine. This showed that, for steady-state conditions, the order of potency for GluA1 block was spermine > NASPM > PhTx-433 > PhTx-74 (**Fig. 2b, c**). The same analysis demonstrated that the potency of each blocker was reduced when GluA1 was co-expressed with TARP γ2 (between 7- and 18-fold reduction; **Fig. 2b, c**).

**Figure 2.**
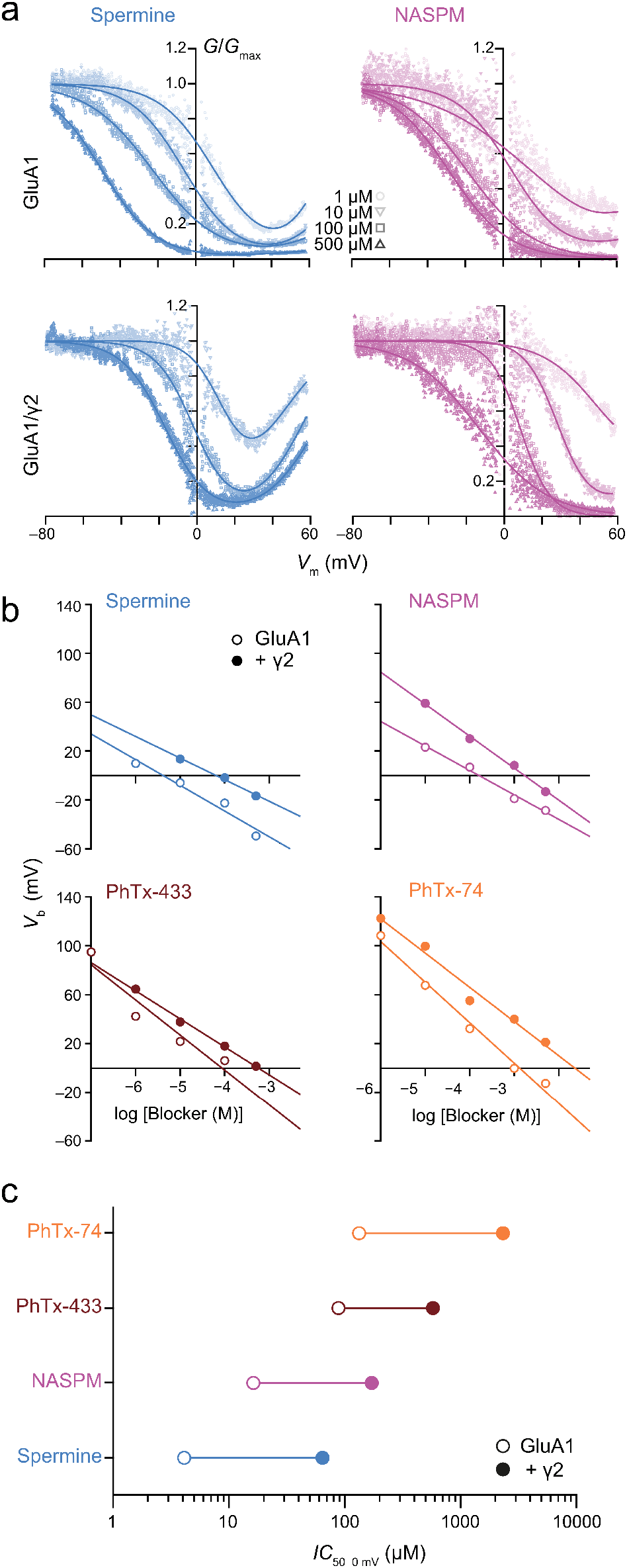
Spermine, NASPM and polyamine toxins display different potencies of GluA1 block that are all decreased by TARP γ2. **a**) Pooled, normalized conductance-voltage relationships (from voltage ramps as in **Figure 1**) for GluA1 and GluA1/γ2 in the presence of spermine (left) or NASPM (right) (*n* = 4-8). Conductance was corrected for the outwardly rectifying response seen in polyamine-free conditions and fitted with single or double Boltzmann relationships (solid lines). **b**) *V*_b_ values from fitted *G-V* relationships of GluA1 and GluA1/γ2 in the presence of varying concentrations of blockers. When *V*_b_ was plotted against log [blocker] a linear relationship could be fitted in all cases, the x-intercept of which gave *IC*_50 0 mV_. **c**) The *IC*_50 0 mV_ values for each polyamine demonstrate relative potency in the order spermine > NASPM > PhTx-433 > PhTx-74, with γ2 co-expression reducing the potency of the blockers by 7-18-fold. **Figure 2—source data 1** *V*_b_ values for GluA1 and GluA1/γ2 obtained with intracellular spermine, NASPM, PhTx-433 or PhTx-74. Fig_2b_source_data.csv **Figure 2—source data 2** *IC*_50 0 mv_ values for GluA1 and GluA1/γ2 obtained with intracellular spermine, NASPM, PhTx-433 or PhTx-74. Fig_2c_source_data.csv

### Onset and recovery of block by NASPM

Next, we examined GluA1/γ2 currents elicited by rapid application of 10 mM glutamate (in the absence of cyclothiazide) and compared the effects of spermine with those of NASPM, the next most potent of the blockers. We first tested the blockers at a concentration of 10 μM, as this allowed us to examine the onset of block. At both positive and negative voltages, glutamate application produced currents that showed a clear peak and rapid desensitization to a steady-state level. However, with intracellular NASPM the steady state-currents at positive voltages were much reduced compared to the corresponding currents at negative voltages (**Fig. 3a**). Comparing the rectification index (RI_+60/-60_) of peak and steady-state currents with spermine or NASPM, two-way ANOVA indicated a main effect of polyamine (*F*_1, 32_ = 22.33, p < 0.0001), a main effect of phase (peak or steady-state; *F*_1, 32_ = 32.72, p < 0.0001) and an interaction between polyamine and phase (*F*_1, 32_ = 15.05, p < 0.0001). The RI_+60/-60_ values obtained with NASPM were less than those obtained with spermine, for both peak and steady-state currents (Peak: 0.37 ± 0.04 *versus* 0.56 ± 0.07, *n* = 11 and 10; unpaired mean difference −0.19 [−0.34, −0.045], p = 0.031, unpaired Welch *t* test.

**Figure 3.**
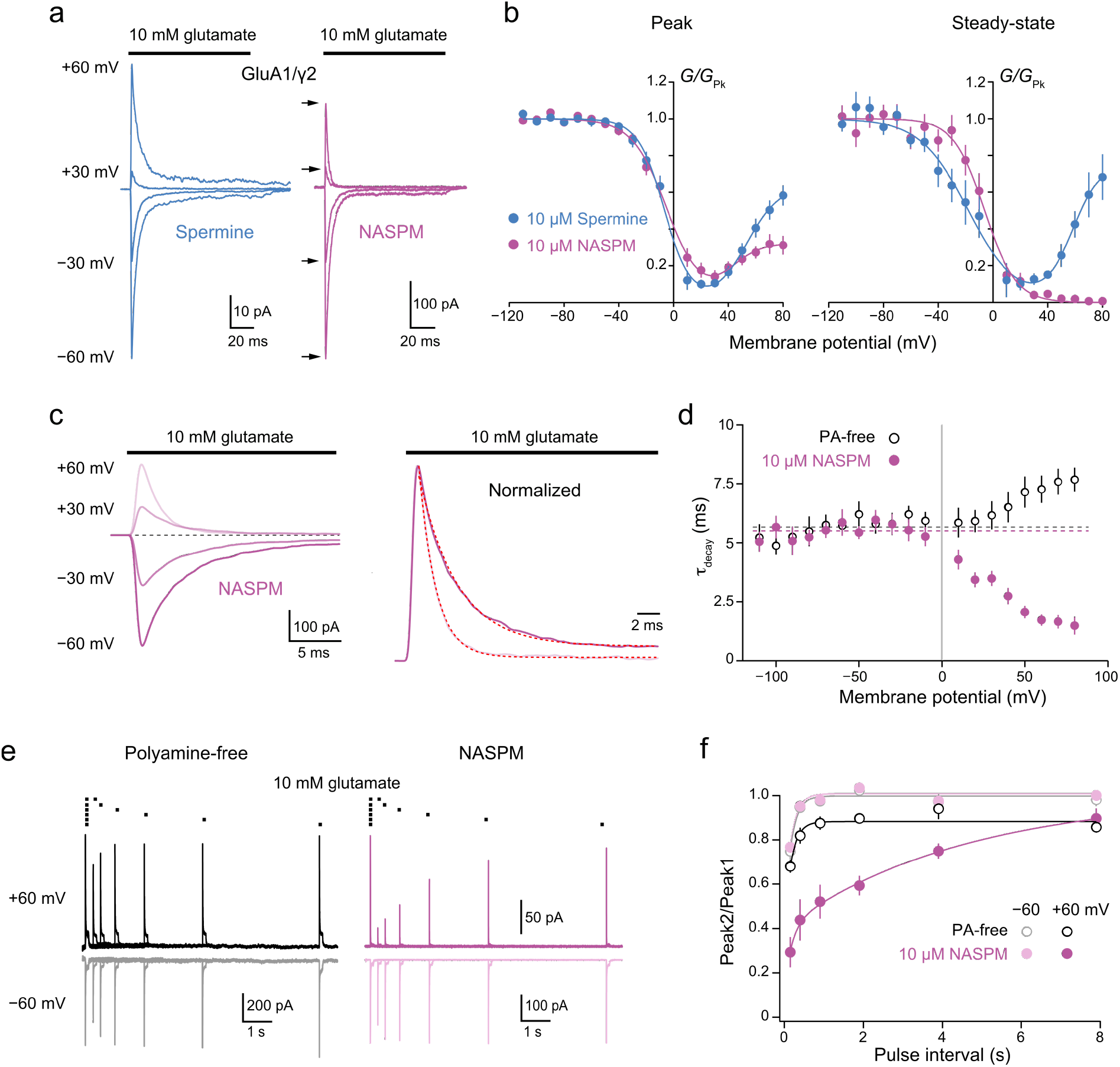
Block of GluA1/γ2 by 10 μM intracellular NASPM shows use-dependence and slow recovery. **a**) Representative GluA1/γ2 currents in the presence of 10 μM intracellular spermine or NASPM activated by fast applications of glutamate (10 mM, 100 ms) at positive and negative potentials. In the presence of NASPM, relative peak outward currents are smaller than those seen with spermine, while steady-state currents are negligible. **b**) Pooled, averaged *G-V* relationships for peak and steady-state currents obtained with spermine (*n* = 9 and 6, respectively) and NASPM (*n* = 11 and 8, respectively). Symbols and error bars indicate mean ± s.e.m. Note the stronger inhibition by NASPM of the steady-state compared with peak currents. **c**) Representative GluA1/γ2 currents with 10 μM intracellular NASPM at the indicated voltages (left) and normalized currents from the same patch at +60 and −60 mV (right). The dashed lines are single exponential fits. **d**) Pooled data showing weighted time constants of decay with polyamine-free (open symbols, *n* = 8) and 10 μM NASPM (*n* = 8) intracellular solutions measured between −110 mV and +80 mV. Symbols and error bars indicate mean ± s.e.m. The kinetics of the negative limb (−110 mV to −10 mV) were voltage and NASPM independent: fitted constant *τ*_decay_ (dashed lines) = 4.79 ms for polyamine free and *τ*_decay_ = 4.84 ms for NASPM. The kinetics of the positive limb (+10 to +80 mV) were markedly accelerated in the presence of intracellular NASPM. **e**) Superimposed GluA1/γ2 currents in the presence or absence of 10 μM NASPM elicited by pairs of glutamate applications at 4, 2, 1, 0.5, 0.25 and 0.125 Hz (10 mM, 100 ms, +/-60 mV). The first responses with NASPM were scaled to those in the polyamine free condition. With NASPM at −60 mV and in the absence of polyamine at both voltages, the second currents broadly recovered to the initial levels. With NASPM at +60 mV however, peak currents recovered much more slowly. **f**) Pooled recovery data recorded in the polyamine-free condition (open symbols, *n* = 9) and with added 10 μM NASPM (filled symbols, *n* = 9). Symbols and error bars indicate mean ± s.e.m. The plots reveal a slow component of recovery present only at +60 mV with NASPM. The biphasic recovery is suggestive of two populations of receptors, those that desensitized before being blocked (fast component), or those that were blocked by NASPM (slow component). **Figure 3–source data 1** Normalised G-V data for peak and steady-state currents evoked by 10 mM glutamate from GluA1/γ2 receptors with intracellular spermine (10 μM) or NASPM (10 μM). Fig_3b_source_data.csv **Figure 3–source data 2** *τ*_decay_ values for currents evoked by 10 mM glutamate from GluA1/γ2 receptors with intracellular NASPM (10 μM) or with polyamine-free intracellular solution at different membrane voltages. Fig_3d_source_data.csv **Figure 3–source data 3** Recovery data (Peak 2/Peak 1) for currents evoked by 10 mM glutamate from GluA1/γ2 receptors with intracellular NASPM (10 μM) or with polyamine-free intracellular solution at +60 and −60 mV. Fig_3f_source_data.csv

Steady-state: 0.04 ± 0.02 *versus* 0.90 ± 0.14, *n* = 8 and 7; −0.86 [−1.12, −0.62], p = 0.00075). This difference was also evident in the *G-V* relationships; those determined from peak responses in the presence of spermine or NASPM were largely overlapping, while those from steady-state responses were markedly different at potentials positive to +40 mV, with a large outward conductance seen in the presence of spermine but not NASPM (**Fig. 3b**).

Intracellular spermine is a weakly permeable open-channel blocker of both CP-AMPARs and kainate receptors (KARs) (Bowie et al., 1998; Brown et al., 2016). However, for GluK2(Q) KARs, permeation of intracellular polyamines has been shown to vary with molecular size, with two molecules of greater width than spermine – *N*-(4-hydroxyphenylpropanoyl)-spermine (HPP-SP) and PhTx-433 – showing little or no detectable relief from block with depolarization (Bahring et al., 1997). Accordingly, NASPM, with its naphthyl headgroup, might also be expected to display limited permeability of CP-AMPAR channels. This could account for the shape of the ramp *I-V* (**Fig. 1c**) and *G-V* plots with NASPM (**Fig. 2a**). Indeed, the pronounced effect of 10 μM intracellular NASPM on steady-state GluA1/γ2 currents at positive potentials (**Fig. 3a**) is also consistent with limited permeation, leading to the accumulation of channel block.

In line with this, we found that the decay of GluA1/γ2 currents recorded in the presence NASPM was strongly voltage-dependent (**Fig. 3a, c**). At negative potentials *τ*_decay_ values were similar in the presence and absence of NASPM. However, at positive potentials (from +10 to +80 mV) the kinetics in the two conditions differed markedly; in the absence of NASPM current decay was slowed at positive potentials, while in the presence of NASPM the decay was progressively accelerated (**Fig. 3d**). In the absence of NASPM, *τ*_decay_ was slower at +60 mV than at −60 mV (7.3 ± 0.6 ms *versus* 5.7 ± 0.3 ms, *n* = 8; paired mean difference 1.55 ms [0.86, 2.62], p = 0.019 paired Welch t-test). By contrast, in the presence of NASPM *τ*_decay_ was markedly faster at +60 mV than at −60 mV (1.7 ± 0.2 ms *versus* 5.6 ± 0.6 ms, *n* = 10; paired mean difference −4.14 ms [−5.44, −3.07], p = 0.00012). Of note, at +60 mV, along with the accelerated decay in the presence of NASPM, we also observed a dramatic slowing of the recovery of peak responses following removal of glutamate (**Fig. 3e**). Currents were elicited by pairs of 100 ms glutamate applications at frequencies from 0.125 to 4 Hz. The peak amplitude of successive responses is normally shaped solely by the kinetics of recovery from desensitization. In the absence of polyamines, a small degree of residual desensitization was apparent at 4 Hz (150 ms interval), but full recovery was seen at all other intervals, at both +60 and −60 mV (**Fig. 3f**). However, in the presence of NASPM, although responses at −60 mV were indistinguishable from those in the absence of polyamines, at +60 mV an additional slow component of recovery (T_rec slow_ 4.9 s; **Fig. 3f**) was present. Taken together, our data suggest that 10 μM intracellular NASPM produces a pronounced, rapid and long-lasting inhibition of CP-AMPARs at positive potentials.

### NASPM can induce complete rectification that is unaffected by auxiliary subunits

We found that with 10 μM NASPM the block of CP-AMPARs was incomplete (**Fig. 3a,b**) and depended on the recent history of the channel (**Fig. 3f**). However, as intracellular polyamines produce both closed- and open-channel block of CP-AMPARs (Bowie et al., 1998; Rozov et al., 1998), we reasoned that a higher concentration of NASPM would block more effectively and cause pronounced rectification, regardless of recent activity. With a higher concentration of NASPM not only would closed-channel block be more favoured, rendering a higher proportion of receptors silent prior to activation, but unblocked closed receptors would more rapidly enter a state of open-channel block once activated at positive potentials. For example, open channel block with 100 μM would be approximately 10 times faster than with 10 μM NASPM and would be expected to rapidly curtail charge transfer.

Importantly, the block of CP-AMPARs by intracellular spermine has been shown to be affected by AMPAR auxiliary proteins. TARPs and CNIHs variously reduce inward rectification (Brown et al., 2018 Cho et al Howe 2007; Coombs et al., 2012; Soto et al., 2007; Soto et al., 2009), while GSG1L increases rectification (McGee et al., 2015). Thus, any experimentally observed change in spermine-induced rectification cannot be unambiguously attributed to a change in CP-AMPAR prevalence alone. Accordingly, we next sought to determine whether NASPM was able to produce complete rectification and whether its action was affected by the presence of auxiliary subunits. To this end, we compared the effects of 100 μM spermine and 100 μM NASPM on *I-V* relationships for GluA1 expressed alone or with γ2, γ7, CNIH2 or GSG1L (**Fig. 4a-c**). With GSG1L, full rectification was seen with both spermine and NASPM (RI_+60/-60_ = 0). For all other combinations, in the presence of spermine RI_+60/-60_ varied (from 0.037 to 0.29) depending on the auxiliary subunit present, but with NASPM rectification was essentially complete in all cases (RI_+60/-60_ 0 to 0.01). Specifically, for GluA1/γ2 RI_+60/-60_ was 0.29 ± 0.04 with spermine and 0.010 ± 0.006 with NASPM (*n* = 6 and 5, respectively; unpaired mean difference −0.28 [−0.34, −0.21], p = 0.00067, unpaired Welch t-test). For GluA1/γ7 RI_+60/-60_ was 0.096 ± 0.024 and 0 ± 0 (*n* = 7 and 5, respectively; unpaired mean difference −0.096 [−0.13, −0.049], p = 0.0073) and for GluA1/CNIH2 the corresponding values were 0.10 ± 0.01 and 0.01 ± 0.01 (*n* = 6 and 5, respectively; unpaired mean difference −0.089 [−0.12, −0.056], p = 0.00095). Thus, unlike 100 μM spermine, 100 μM NASPM produces a near total block of outward CP-AMPAR-mediated currents, independent of auxiliary subunit association.

**Figure 4.**
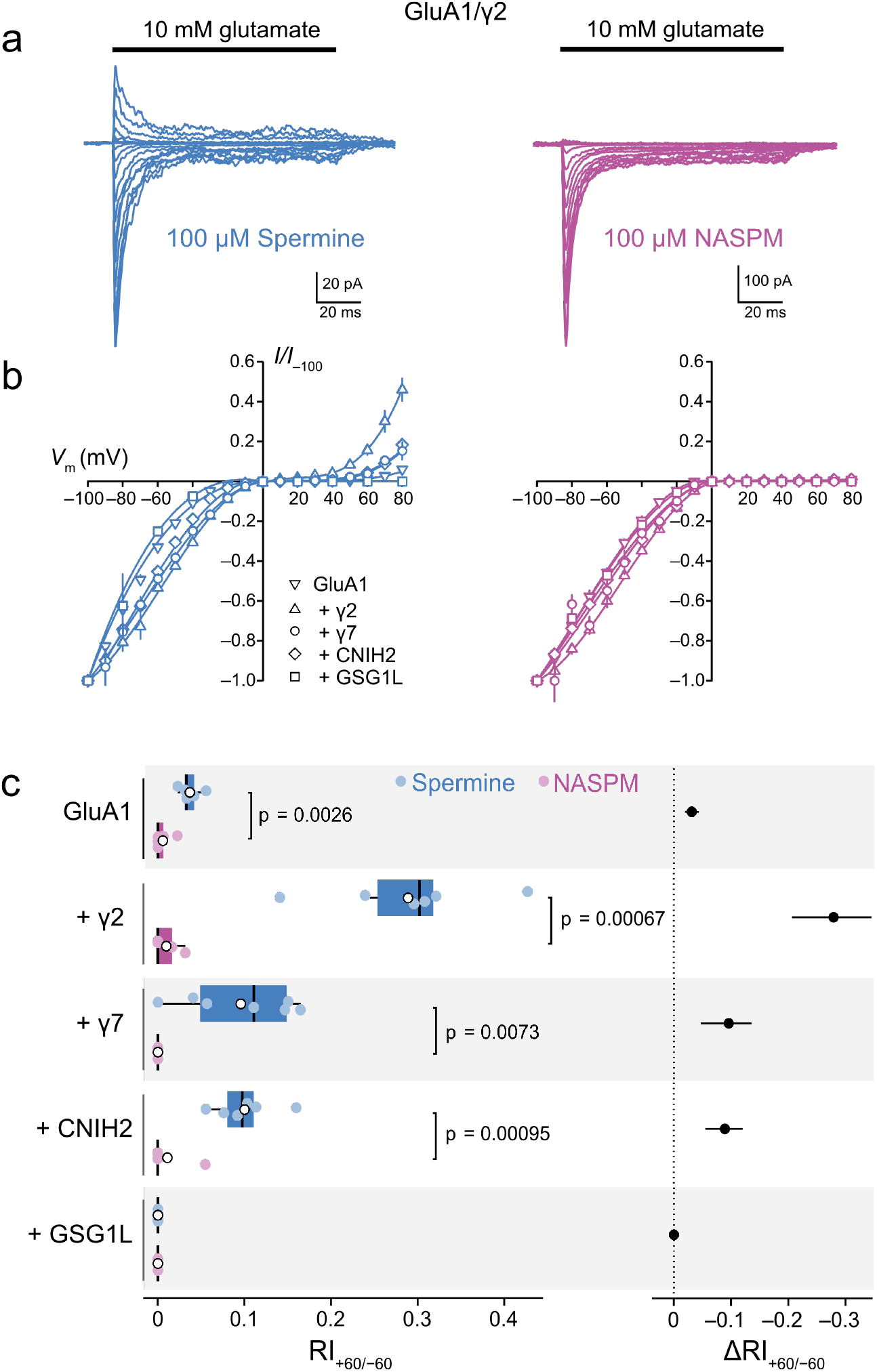
Rectification in the presence of 100 μM intracellular NASPM is complete and unaffected by auxiliary proteins. **a**) Representative GluA1/γ2 currents activated by fast applications of glutamate to outside-out patches excised from transfected HEK293 cells (10 mM, 100 ms, −110 to +80 mV). The intracellular solution was supplemented with 100 μM spermine (left) or NASPM (right). **b**) Mean *I-V* relationships for GluA1 expressed alone or with γ2, γ7, CNIH2 or GSG1L with 100 μM spermine (left) or NASPM (right). Symbols and error bars indicate mean ± s.e.m. Negligible outward currents are seen for any combination in the presence of NASPM. **c**) Pooled rectification data (RI_+60/-60_) for GluA1 combinations with 100 μM spermine or NASPM (left). Box-and-whisker plots indicate the median (black line), the first and third quartiles (Q1 and Q3; 25-75th percentiles) (box), and the range min(x[x > Q1 + 1.5 * interquartile range]) to max(x[x < Q3 + 1.5 * inter-quartile range]) (whiskers). Indicated p values are from Welch two-sample *t* tests. Difference plots showing the shift in RI_+60/-60_ in the presence of NASPM compared with spermine (right). Symbols show mean unpaired differences, and the error bars indicate the bootstrapped 95% confidence intervals. **Figure 4–source data 1** Normalised *I-V* data for peak currents evoked by 10 mM glutamate from GluA1, GluA1/γ2, GluA1/γ7, GluA1/CNIH2 and GluA1/GSG1L receptors with intracellular spermine (100 μM) or NASPM (100 μM). Fig_4b_source_data.csv **Figure 4–source data 2** RI_+60/-60_ values for peak currents evoked by 10 mM glutamate from GluA1, GluA1/γ2, GluA1/γ7, GluA1/CNIH2 and GluA1/GSG1L receptors with intracellular spermine (100 μM) or NASPM (100 μM). Fig_4c_source_data.csv

### NASPM fully blocks CP-AMPAR-mediated mEPSCs at positive voltages

To determine whether 100 μM intracellular NASPM would cause complete rectification of CP-AMPAR-mediated synaptic currents, we recorded mEPSCs from stellate cells in cerebellar cultures prepared from GluA2^-/-^ (*Gria2^tm1Rod^*) mice (Jia et al., 1996). Functional studies have shown these molecular layer interneurons normally possess both CI- and CP-AMPARs (Liu and Cull-Candy, 2002; Liu and Cull-Candy, 2000), formed from GluA2, −3 and −4 subunits (Yamasaki et al., 2011) that are expressed along with the TARPs γ2 and γ7 (Bats et al., 2012; Fukaya et al., 2005; Yamazaki et al., 2010). Thus, in stellate cells from GluA2^-/-^ mice it is expected that AMPAR-mediated EPSCs will be mediated by TARP-associated GluA2-lacking CP-AMPARs. With 100 μM spermine added to the intracellular solution mEPSCs were detected in stellate cells from GluA2^-/-^ mice at both negative and positive voltages (**Fig. 5a, c**). At −60 mV the mean absolute amplitude of the averaged mEPSCs was 39 ± 5 pA, the 20-80% rise time was 0.21 ± 0.02 ms and the *τ*_w,decay_ was 1.14 ± 0.09 ms (*n* = 6). At +60 mV the corresponding measures were 22 ± 2 pA, 0.23 ± 0.03 ms and 1.58 ± 0.19 ms. In each cell, fewer events were detected at +60 mV that at −60 mV (the mean frequency was 2.3 ± 1.6 Hz at +60 mV and 7.7 ± 5.0 Hz at −60 mV). This is consistent with the anticipated blocking action of spermine making a proportion of mEPSCs undetectable at positive potentials (McGee et al., 2015). To take into account the apparent change in frequency when assessing the degree of rectification, the rectification index (RI_+60/-60_) was calculated from the mean absolute amplitude of detected events multiplied by the frequency at each voltage (see **Materials and methods**). With spermine, the calculated RI_+60/-60_ was 0.19 ± 0.04. When we made recordings with 100 μM intracellular NASPM, mEPSCs were detectable at negative but not positive voltages (**Fig. 5b, c**). At −60 mV the mean absolute amplitude of the averaged mEPSCs was 36 ± 6 pA, the 20-80% rise time was 0.26 ± 0.02 ms and the *τ*_w,decay_ was 1.45 ± 0.17 ms (*n* = 6). At −60 mV the average mEPSC frequency was 14.2 ± 8.2 Hz (*n* = 6), but at +60 mV no events were seen (RI_+60/-60_ = 0; **Fig. 5d**). Thus, even when synapses contained exclusively GluA2-lacking AMPARs, spermine did not fully block outward currents. By contrast, NASPM produced full inward rectification, providing an unambiguous read-out of a pure population of CP-AMPARs.

**Figure 5.**
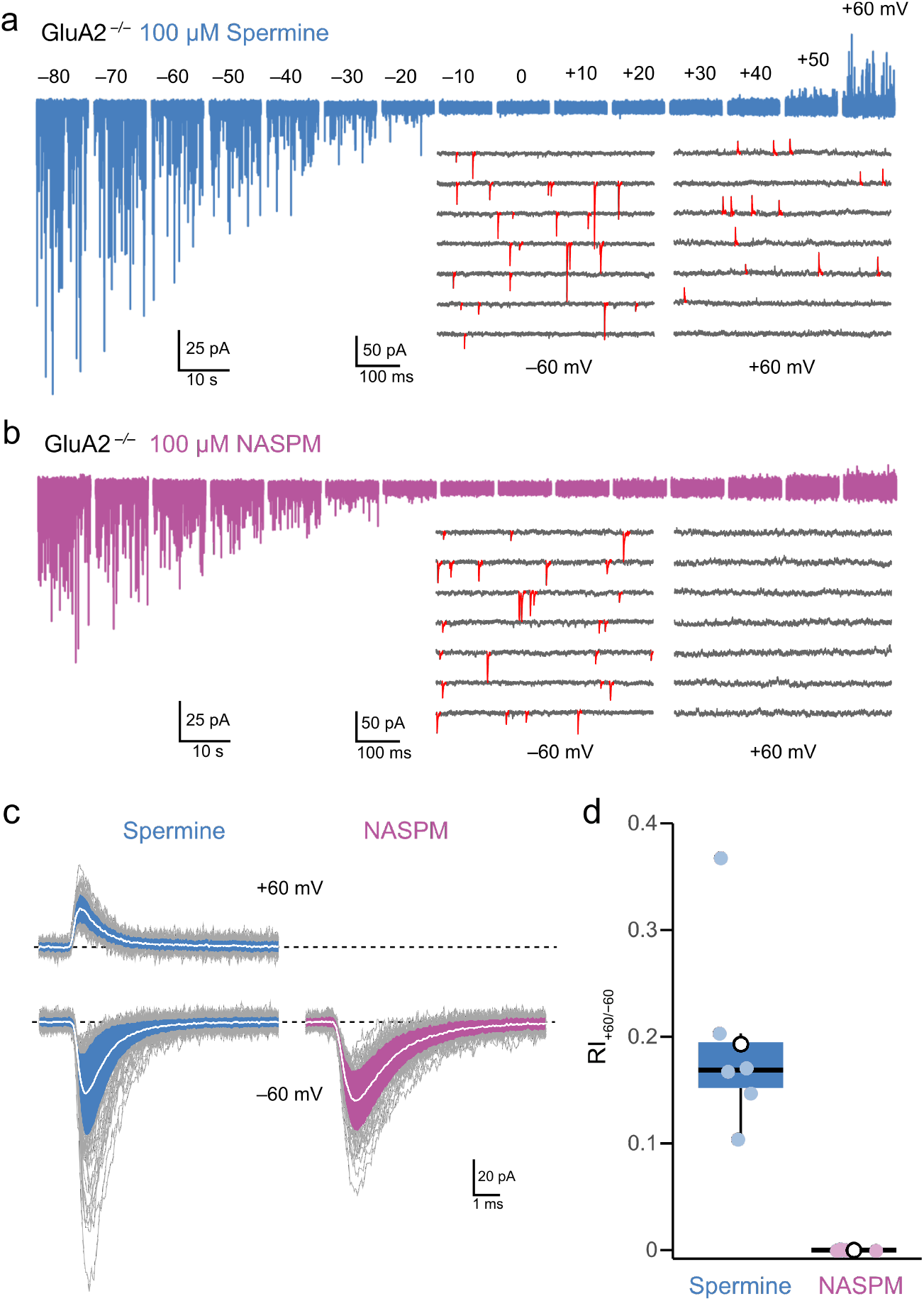
Intracellular NASPM (100 μM) produces full rectification of CP-AMPAR-mediated mEPSCs. **a**) Representative whole cell recording from a cultured GluA2^-/-^ cerebellar stellate cell held at voltages between −80 and +60 mV with 100 μM intracellular spermine. The inset shows expanded sweeps at −60 and +60 mV, with mEPSCs indicated in red. Note the presence of mEPSCs at both voltages. **b**) Same as a, but for a cell with 100 μM intracellular NASPM. The inset shows that mEPSCs (red) were present at −60 mV but not at +60 mV. **c**) Representative averaged mEPSCs from a cell recorded with intracellular spermine (left) and from a cell recorded with intracellular NASPM (right). Individual mEPSCs are shown in gray, the averaged mEPSCs are shown in white, superimposed on filled areas indicating the standard deviation of each recording. **d**) Box-and-whisker plot (as described in Fig. 4c) of pooled RI_+60/-60_ data. **Figure 5–source data 1** RI_+60/-60_ values for cerebellar stellate cell mEPSCs recorded with intracellular spermine (100 μM) or NASPM (100 μM). Fig_5d_source_data.csv

## Discussion

Our results demonstrate that when included in the intracellular recording solution the exogenous polyamine toxins PhTx-433 and PhTx-74 and the toxin analogue NASPM cause selective, voltage-dependent block of GluA2-lacking CP-AMPARs. With GluA1 AMPARs the order of potency is spermine > NASPM > PhTx-433 > PhTx-74. As with spermine (Soto et al., 2007), the potency of the blockers is reduced when the receptors are associated with the TARP γ2. However, in contrast with spermine, which is currently widely used as means of identifying CP-AMPARs, NASPM can produce a complete block of outward currents through CP-AMPARs. Furthermore, the block remains complete when CP-AMPARs are co-assembled with auxiliary subunits. These characteristics make the use of intracellular NASPM a valuable new approach to disclose the presence of functional CP-AMPARs. Indeed, given that spermine is often added to the pipette solutions when recording EPSCs and other AMPAR-mediated currents solely to allow rectification to be used as a proxy for the presence of CP-AMPARs, a strong case can be made that use of 100 μM intracellular NASPM should become the new standard approach in the field.

### The mechanism of polyamine toxin block

The *G-V* relationships we obtained over a range of concentrations of intracellular blockers were fitted by single (block) or double (block and permeation) Boltzmann functions (Panchenko et al., 1999). As spermine can permeate the channel (Brown et al., 2018), double Boltzmann functions were required to adequately describe the full *G-V* relationships for all concentrations across all protocols. Double Boltzmann functions were also needed to describe both ramp and peak responses recorded with 1 or 10 μM intracellular NASPM, where there was a small decrease in current block at potentials more positive than +40 mV. This possibly reflects a degree of blocker permeation, as suggested for extracellularly applied NASPM or adamantane derivates at hyperpolarized voltages (Tikhonov et al., 2000; Twomey et al., 2018). However, with 100 μM or greater intracellular NASPM, and with all concentrations of PhTx-433 and PhTx-74 examined, outward currents were not detected at positive voltages (giving full inward rectification) and single Boltzmann fits were adequate.

Recent cryo-EM structures of activated GluA2/γ2 AMPARs blocked with NASPM show the hydrophobic naphthalene moiety sitting within the electroneutral central vestibule and the polyamine tail extending into the electronegative selectivity filter (Twomey et al., 2018). This is in accord with predictions from molecular modelling of the block by various philanthotoxins and adamantane derivatives (Andersen et al., 2006; Tikhonov, 2007) and suggests that the majority of the observed receptors were blocked from the ‘outside’. Our data show, unsurprisingly, that the voltage-dependence of block by intracellular NASPM is the inverse of that seen with extracellular NASPM (Koike et al., 1997; Twomey et al., 2018). Given that extracellular NASPM block is removed at positive potentials, this suggests that intracellularly applied NASPM occupies the pore in an ‘inverted’ orientation, with its polyamine tail in the selectivity filter, but with its bulky head group protruding to the intracellular side of the channel. The differential action of intracellular and extracellular PhTx-74 on heteromeric CI-AMPARs is consistent with this interpretation. These CI-AMPARs are blocked by extracellular PhTx-74 (Jackson et al., 2011; Nilsen and England, 2007; Poulsen et al., 2014) but are unaffected when this molecule is applied from the inside (Fig. 1). This route-specific differential effect of PhTx-74 on CI-AMPARs may arise because its polyamine tail contains only three of the four amine groups found in the other toxins, which leads to an asymmetric distribution. When PhTx-74 is applied from the outside, the electrostatic repulsion mediated by the edited Q/R site may be limited due to the missing amine. When present inside the cell, however, the proposed ‘inverted’ toxin orientation would maintain Q/R site repulsion through the intact terminal amine groups.

Intriguingly, in the cryo-EM study of Twomey et al (2018), the purified receptors (in effect at 0 mV in a cell-free environment) were incubated with NASPM for an extended period and glutamate was subsequently applied immediately prior to freezing. Thus, while the channel was closed, NASPM could only access the selectivity filter from the ‘inside’. Despite this, the finally determined orientation of the toxin suggests that once the channels were opened any closed- or open-channel block by ‘intracellular’ NASPM was rapidly replaced with open-channel block by ‘extracellular’ NASPM. Our estimate of the concentration of intracellular NASPM required to block 50% of the channels at 0 mV (160 μM for GluA1/γ2) reveals a 4-fold lower potency than found for extracellular NASPM (35 μM for GluA2_Q_-STZ; Twomey et al., 2018). This difference could reflect, for extracellular block, a permissive environment for the headgroup in the central vestibule of the channel or a different alignment of amines within the polyamine tail with anchoring points in the electronegative selectivity filter (Twomey et al., 2018). Alternatively, when entering from the intracellular side, the NASPM head group may be sterically hindered by the Q/R +4 aspartate residues, so decreasing the affinity in this orientation.

We have shown previously that with GluA4, γ2 reduces spermine potency by around 20-fold (Soto et al., 2007). This is thought to result from increased spermine permeation (Brown et al., 2018). Here, we found that co-expression of γ2 with GluA1 reduced the potency of all intracellular blockers (7 to 18-fold). Unlike spermine, however, NASPM, PhTx-74 and PhTx-433 show either no permeation or limited permeation (at low concentrations), so the γ2 alteration of *G-V* relationships likely reflects instead the facilitation of blocker unbinding from the open pore to the intracellular side. Whether this arises from TARP-dependent reshaping of the selectivity filter, as suggested for auxiliary protein modulation of intracellular spermine block (Brown et al., 2018; Soto et al., 2014), remains to be determined.

### Advantages of intracellular NASPM for assessing CP-AMPAR contribution

Although intracellular spermine is widely used in voltage-clamp recordings to assess the contribution of CP-AMPARs to the recorded currents, intracellular NASPM offers two key advantages. First, it can produce a selective and complete block of CP-AMPAR-mediated outward current. Second, this block is unaffected by the presence of auxiliary proteins. Together, these characteristics eliminate the difficulties of interpretation that can arise when using spermine.

Rectification of AMPAR-mediated currents is commonly quantified using a rectification index (RI). This is frequently the ratio of conductance or current amplitudes at +40 (or +60) and −60 mV (e.g. Castilho et al., 2015; Fortin et al., 2010; Mahanty and Sah, 1998; Ozawa et al., 1991; Studniarczyk et al., 2013; Sullivan et al., 2017), but ratios at various arbitrary voltages or over various voltage ranges have been used. Because *I-V* relationships of CP-AMPAR-mediated currents recorded with intracellular spermine are typically bi-rectifying, the choice of these voltages (particularly the positive voltage) can influence the absolute value of RI. With intracellular NASPM, however, outward current through CP-AMPARs is fully blocked at all positive voltages, potentially allowing greater sensitivity to detect the contribution from a small population of CP-AMPARs. Concomitantly, this full block of CP-AMPARs at positive potentials will allow any limited contribution of CI-AMPARs to be more effectively assessed, as recordings may be made at more depolarized potentials with increased driving force reducing the chance that CI-AMPAR-mediated currents are missed. These considerations apply under all conditions but particularly so when the receptors are associated with TARPs or CNIHs which partially relieve spermine-dependent rectification (Brown et al., 2018; Cho et al., 2007; Coombs et al., 2012; Soto et al., 2007; Soto et al., 2009). With intracellular spermine, an intermediate level of rectification is difficult to interpret, as it could reflect the presence of a mixture of CI- and CP-AMPARs or the presence of CP-AMPARs with various auxiliary proteins. With NASPM, however, incomplete rectification can arise only if there is a genuine contribution of CI-AMPARs.

Whilst a principal advantage of NASPM is its insensitivity to the presence of auxiliary subunits, this means that it cannot discriminate between different CP-AMPAR subtypes with potentially different auxiliary subunit content. By contrast, we have previously used the precise degree of rectification seen with spermine to infer the association of auxiliary subunits with native CP-AMPARs (Soto et al., 2007; Soto et al., 2009; Studniarczyk et al., 2013). Thus, we suggest that intracellular NASPM will prove most informative when used in parallel with intracellular spermine. For example, as shown in **Fig. 5**, when recorded with NASPM, the mEPSCs from GluA2^-/-^ stellate cells have an RI of 0, which indicates that the mEPSCs are generated entirely by CP-AMPARs. If this observation came from an experiment in which the nature of the GluA2 expression was not known, it would allow a clearer interpretation of any RI obtained in parallel experiments with spermine, as the contribution of CI-AMPARs could be excluded. Comparison of this spermine RI with corresponding measures from recombinant receptors with and without different auxiliary subunits (as shown in **Fig. 4**) may allow one to infer the presence or absence of specific auxiliary subunits. Without the use of NASPM, information from other measures, such as the channel conductance estimated using nonstationary fluctuation analysis (Bats et al., 2012; Studniarczyk et al., 2013), is needed to make such inferences.

The extracellular application of cationic blockers, such as HPP-SP, NASPM, PhTx-433 or IEM-1460, has been widely used to reveal the presence and roles of CP-AMPARs in neurons, with the advantage that recordings can be made at a fixed negative voltage and the block can be reversible. However, those extracellular blockers that have been examined on CP-AMPAR-mediated synaptic responses in tissue from GluA2^-/-^ mice – HPP-SP, PhTx-433 and IEM-1460 – produce an incomplete (~60-80%) block, even when used at relatively high concentrations (10 or 100 μM) (Adesnik and Nicoll, 2007; Gray et al., 2007; Jackson and Nicoll, 2011b; Mainen et al., 1998; Sara et al., 2011). In this respect it is revealing that a low concentration (100 nM) of PhTx-433, that is capable of producing near complete steady-state block of recombinant GluA2-lacking receptors, has only a minimal effect on GluA2-lacking synaptic responses (Jackson and Nicoll, 2011b). This difference in efficacy reflects the different nature of glutamate exposure and the use-dependent nature of the block. This also means that, when used to examine the properties of synaptic responses, the effects can be frequency-dependent and exhibit a slow onset (Lujan et al., 2019; Mainen et al., 1998; Zaitsev et al., 2011). Finally, of course, extracellularly applied drugs have the opportunity to block CP-AMPARs throughout the tissue, not only those in the recorded cell. This can potentially affect glutamate release, either directly or indirectly.

To conclude, our recordings of macroscopic currents from heterologously expressed recombinant AMPARs and mEPSCs from dissociated neurons have demonstrated the value of intracellular NASPM as a tool for the functional study of AMPARs. We suggest that for any tissue, cell type or recording scenario where spermine-dependent rectification may be used to assess CP-AMPAR expression, intracellular NASPM offers a clear methodological advance, enabling a straightforward, cell-specific readout of CP-AMPAR presence and the ability to unambiguously assess changes in their prevalence.

## Materials and methods

### Materials

Common laboratory chemicals were obtained from Sigma-Aldrich (Merck Life Science UK Limited, Gillingham, UK), as were penicillin, streptomycin, transferrin, insulin, spermine tetrahydrochloride and PhTx-433. Cyclothiazide, TTX, d-AP5, SR-95531 and PhTx-74 were from Tocris Bioscience (Bio-Techne Ltd, Abingdon, UK). Lipofectamine 2000 and gentamicin were from Invitrogen (Thermo Fisher Scientific, Waltham, MA USA). Basal Medium Eagle (BME), Eagles Minimum Essential Medium (MEM), Dulbecco’s Modified Eagle Medium (DMEM) and fetal bovine serum (FBS) were from Gibco (Thermo Fisher Scientific, Waltham, MA USA). NASPM was from Hello Bio (Bristol, UK).

### Mice

*Gria2*-deficient (GluA2^-/-^) mice were bred from heterozygous parents (*Gria2^tm1Rod^*, MGI: 1857436) (Jia et al., 1996). Mice were group housed in standard cages and maintained under controlled conditions (temperature 20 ± 2°C; 12 h light-dark cycle). Food and water were provided *ad libitum*. Both male and female mice were used for generating primary neuronal cultures. To increase likelihood of homozygotes surviving, at P1 the litter numbers were typically reduced to six, by culling the largest pups (those presumed to be wild-type or heterozygous). The genotypes of the remaining pups were subsequently confirmed using PCR analysis using the following primers: oIMR6780 GGTTGGTCACTCACCTGCTT (wild type); oIMR6781 TCGCCCATTTTCCCATATAC (common) and oIMR8444 GCCTGAAGAACGAGATCAGC (mutant). All procedures for the care and treatment of mice were in accordance with the Animals (Scientific Procedures) Act 1986.

### Heterologous expression

The flip splice variant of rat GluA1, the Q/R and R/G edited flip splice variant of rat GluA2, rat γ2, CNIH2 and GSG1L, and human γ7 were expressed from the pIRES2-EGFP vector. HEK293T/17 cells (ATCC; Manassas, VA, USA) were grown in DMEM supplemented with 10% fetal bovine serum (FBS), 100 U/ml penicillin, 0.1 mg/ml streptomycin at 37°C, 5% CO_2_ and maintained according to standard protocols. Transient transfection was performed using Lipofectamine 2000 according to the manufacturer’s instructions. Heteromeric GluA1/2 receptors were expressed using a cDNA ratio of 1:2 GluA1:GluA2. AMPAR/auxiliary subunit combinations had a cDNA ratio of 1:2 GluA1:auxiliary subunit. Cells were split 12–24 h after transfection and plated on poly-L-lysine coated glass coverslips. Electrophysiological recordings were performed 18–48 h later.

### Outside-out patch recordings

Patch-clamp electrodes were pulled from borosilicate glass (1.5 mm o.d., 0.86 mm i.d.; Harvard Apparatus, Cambridge, UK) and fire polished to a final resistance of 6–10 MΩ. Voltage ramp experiments were performed using outside-out patches. The ‘external’ solution contained: 145 mM NaCl, 2.5 mM KCl, 1 mM CaCl_2_, 1 mM MgCl_2_ and 10 mM HEPES (pH 7.3). For steady-state ramp responses this solution was supplemented with 300 μM glutamate and 50 μM cyclothiazide. The ‘internal’ solution contained 125 mM CsCl, 2.5 mM NaCl, 1 mM CsEGTA, 10 mM HEPES and 20 mM Na_2_ATP (pH 7.3 with CsOH) and was supplemented with the indicated concentrations of spermine tetrahydrochloride, NASPM, PhTx-433 or PhTx-74. For fast-application experiments, the ‘external’ solution contained 145 mM NaCl, 2.5 mM KCl, 2 mM CaCl_2_, 1 mM MgCl_2_ and 10 mM HEPES (pH 7.3) while the ‘internal’ solution contained (in mM): 140 mM CsCl, 10 mM HEPES, 5 mM EGTA, 2 mM MgATP, 0.5 mM CaCl_2_ and 4 mM NaCl, supplemented with 10 or 100 μM spermine or NASPM as indicated. Recordings were made at 22–25°C using an Axopatch 200A amplifier (Molecular Devices, USA). Voltage ramp currents were low-pass filtered at 500 Hz and digitized at 2 kHz using an NI USB-6341 (National Instruments) interface with WINWCP (Strathclyde Electrophysiology Software; http://spider.science.strath.ac.uk/sipbs/software_ses.htm). Recordings of responses to fast glutamate applications were low pass filtered at 10 kHz and digitized at 20 kHz.

Voltage ramps (either −80 mV to +60 mV, −100 mV to +100 mV or−-100 mV to +130 mV) were delivered at 100 mV/s. For each experiment, ramps in control or glutamate plus cyclothiazide solutions were interleaved to allow for leak subtraction. Rapid agonist application was achieved by switching between continuously flowing solutions, as described previously (Soto et al., 2014). Solution exchange was achieved by moving an application tool – made from custom triple-barrelled glass (VitroCom, Mountain Lakes, NJ, USA) – mounted on a piezoelectric translator (PI (Physik Instrumente) Ltd, Bedford, UK). At the end of each experiment, the adequacy of the solution exchange was tested by destroying the patch and measuring the liquid-junction current at the open pipette (10–90% rise time typically 150–300 μs).

### Neuronal primary culture

Primary cultures were prepared from the cerebella of P7 mice as previously described (McGee et al., 2015). After dissociation, the cell suspension was transfected with a Homer 1c::TdimerDsRed plasmid (Bats et al., 2007) by electroporation using the Amaxa Nucleofector 2b device and Amaxa mouse neuron nucleofector kit (Lonza). Neurons were then plated on poly-L-lysine-coated glass coverslips and grown at 37°C in a humidified 5% CO_2_-containing atmosphere in BME supplemented with KCl (25 mM final concentration), 20 μg/ml gentamicin, 2 mM L-glutamine and 10% v/v fetal bovine serum. After 3 days *in vitro* the growing medium was replaced with MEM supplemented with 5 mg/ml glucose, 2 mM glutamine, 20 μg/ml gentamicin, 0.1 mg/ml transferrin and 0.025 mg/ml insulin. Recordings were performed after 8-11 DIV.

### mEPSC recordings

Cells were visualized using an inverted microscope (IX71; Olympus) equipped with a 40×/0.9 NA objective (Olympus) and excitation and emission filters (Chroma Technology HQ540/40x and HQ600/50m) to enable DsRed visualization. Stellate cells were distinguished from granule cells by their larger soma and punctate DsRed labelling of their soma and dendrites that differed from the sparse labelling of granule cells, usually in more distal regions of their dendrites.

The extracellular solution, adjusted to pH 7.3 with NaOH, contained: 145 mM NaCl, 2.5 mM KCl, 2 mM CaCl_2_, 1 mM MgCl_2_, 10 mM glucose, and 10 mM HEPES. To this we added 0.5 μM TTX, 20 μM d-AP5, and 20 μM SR-95531 [2-(3-carboxypropyl)-3-amino-6-(4-methoxyphenyl)pyridazinium bromide] to block voltage-gated sodium channels, NMDA and GABAa receptors, respectively. The internal solution, adjusted to pH 7.4 with CsOH, contained: 140 mM CsCl, 10 mM HEPES, 5 mM EGTA, 2 mM MgATP, 0.5 mM CaCl_2_, 4 mM NaCl, and either 100 μM spermine or 100 μM NASPM. Recordings were performed using an Axopatch 700B amplifier, low pass filtered at 6 kHz, and digitized at 25 kHz using a USB-6341 interface and Igor Pro 6 (v6.35, Wavemetrics, Lake Oswego, Oregon, USA) with NeuroMatic v2.8 and NClamp (http://www.neuromatic.thinkrandom.com) (Rothman and Silver, 2018). Patch-clamp electrodes were pulled from borosilicate glass (as for outside-out patch recordings) and had a resistance of 5-7 MΩ. *R*_series_ was not compensated and was <18 MΩ. *R*_input_ was > 45 × *R*_series_. Cells were exposed for 2 min to 200 μM LaCl_3_ prior to data acquisition to increase mEPSC frequency (Chung et al., 2008). Membrane potential was stepped in 10 mV increments from −80 to +60 mV and then from +60 to −80 mV. For calculation of mEPSC RI_+60/-60_, where both ascending and descending runs were completed, the data were pooled.

### Data analysis

#### Analysis of HEK cell recordings

Records were analyzed using Igor Pro with NeuroMatic. To quantify the rectification seen with ramp *I-V* relationships, the rectification index (RI_+60/-60_) was calculated as the ratio of the current at +60 mV and −60 mV (average of 15 data points spanning each voltage). To generate conductance-voltage (*G-V*) curves the reversal potential for each leak-subtracted *I-V* curve was calculated to ascertain the driving force. The resultant data were normalized, averaged, then converted to conductances. To account for the polyamine-independent outward rectification of AMPARs, the conductance values were divided by those obtained in the polyamine-free condition. For currents that displayed inward rectification only, *G-V* curves were fitted with the Boltzmann equation:

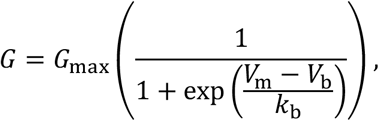

where *G*_max_ is the conductance at a sufficiently hyperpolarized potential to produce full relief of polyamine block, *V*_m_ is the membrane potential, *V*_b_ is the potential at which 50% of block occurs, and *k*_b_ is a slope factor describing the voltage dependence of block (the membrane potential shift necessary to cause an e-fold change in conductance). For currents that displayed double rectification, *G-V* curves were fitted with a double Boltzmann equation which contains equivalent terms for voltage-dependent permeation (p) (Panchenko et al., 1999):

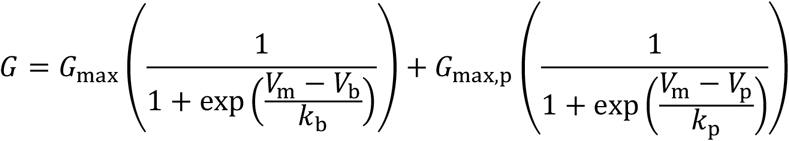

Plots of *V*_b_ against the log of the polyamine concentration were fitted with a linear function, the x-axis intercepts giving the voltage-independent affinity (*IC*_50, 0 mv_).

Current decays following fast applications of glutamate (10 mM, 100 ms) at positive and negative potentials were described by single or double exponential fits. In the latter case, the weighted time constant of decay (*τ*_w,decay_) was calculated according to:

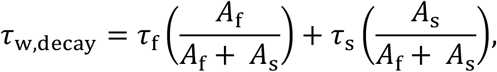

where *A_f_* and *τ*_f_ are the amplitude and time constant of the fast component and *A*_s_ and *τ*_s_ are the amplitude and time constant of the slow component. For each condition, values of *τ*_decay_ (from single exponential fits) and *τ*_w_,_decay_ (from double exponential fits) were pooled.

Double pulse experiments (100 ms glutamate applications at intervals of 150 ms to 7.9 s (4 to 0.125 Hz) were used to assess the recovery of peak responses following removal of glutamate. As expected, some residual desensitization was apparent at the shortest of the intervals in all conditions (Coombs et al., 2017). Thus, the magnitude of the second pulse reflected recovery from desensitization and, at +60 mV with 10 μM NASPM, the relief of NASPM block. In the latter case, the recovery of the second peak was fitted with a biexponential function, as above.

#### mEPSC analysis

mEPSCs were detected using an amplitude threshold crossing method based on the algorithm of (Kudoh and Taguchi, 2002) (NeuroMatic). The standard deviation of the background noise at +60 mV (range 2.4 to 6.4 pA; 3.5 ± 0.4 pA, *n* = 12) was determined by fitting a single-sided gaussian to an all-point histogram from a 500 ms stretch of record. For each cell the same detection threshold (2-3x the standard deviation at +60 mV) was used at both −60 and +60 mV. At each voltage, the mEPSC frequency was determined as the total number of mEPSCs detected/record length, and a mean mEPSC waveform was constructed from those events that displayed a monotonic rise and an uncontaminated decay. One cell recorded with intracellular spermine was excluded from the analysis as <10 events were detected during 40 s recording at −60 mV. For each of the 12 other cells, detected events were aligned on their rising phase before averaging. The rectification index was then calculated as

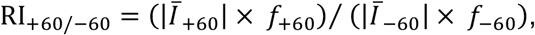

where *Ī* is the amplitude of the mean mEPSC and *f* is the frequency of mEPSCs at the indicated voltage. For each averaged mEPSC we determined the 20–80% rise time and fitted the decay with a double exponential to obtain *τ*_w,decay_ (as described for HEK cell agonist-evoked currents).

### Data presentation and statistics

Summary data are presented in the text as mean ± s.e.m. from *n* measures, where *n* represents the number of biological replicates (number of cells or patches in each data set, from 1-5 separate cultures/transfections). Estimates of paired or unpaired mean differences and bias-corrected and accelerated 95% confidence intervals from bootstrap resampling (Weiss, 2016) are presented as: effect size [lower bound, upper bound]. Error bars on graphs indicate s.e.m. or, in **Fig 4c**, the 95% CI. Box-and-whisker plots (**Fig. 4c** and **Fig. 5c**) indicate the median (black line), the 25-75th percentiles (box), and the 10-90th percentiles (whiskers); filled circles are data from individual patches/cells and open circles indicate means. Comparisons involving two data sets were performed using a two-sided Welch two-sample *t* test that does not assume equal variance (normality was not tested statistically but gauged from density histograms and/or quantile-quantile plots). Analyses involving data from three or more groups were performed using two-way analysis of variance (Welch heteroscedastic *F* test) followed by pairwise comparisons using two-sided Welch two-sample *t* tests. Exact p values are presented to two significant figures, except when p < 0.0001. Statistical tests were performed using R (version 3.6.0, the R Foundation for Statistical Computing, http://www.r-project.org/) and R Studio (version 1.3.1056, RStudio). No statistical test was used to predetermine sample sizes. No randomization was used.

## Acknowledgements

This work was supported by an MRC Programme Grant (MR/T002506/1) to MF and SGCC.

## References

Adesnik H and Nicoll RA (2007) Conservation of glutamate receptor 2-containing AMPA receptors during long-term potentiation. J Neurosci 27:4598–4602. doi: 10.1523/JNEUROSCI.0325-07.2007.

Andersen TF, Tikhonov DB, Bolcho U, Bolshakov K, Nelson JK, Pluteanu F, Mellor IR, Egebjerg J and Stromgaard K (2006) Uncompetitive antagonism of AMPA receptors: Mechanistic insights from studies of polyamine toxin derivatives. J Med Chem 49:5414–5423. doi: 10.1021/jm060606j.

Bahring R, Bowie D, Benveniste M and Mayer ML (1997) Permeation and block of rat GluR6 glutamate receptor channels by internal and external polyamines. J Physiol 502:575–589. doi: 10.1111/j.1469-7793.1997.575bj.x.

Baranovic J and Plested AJ (2016) How to build the fastest receptor on earth. Biol Chem 397:195–205. doi: 10.1515/hsz-2015-0182.

Bats C, Groc L and Choquet D (2007) The interaction between Stargazin and PSD-95 regulates AMPA receptor surface trafficking. Neuron 53:719–734. doi: 10.1016/j.neuron.2007.01.030.

Bats C, Soto D, Studniarczyk D, Farrant M and Cull-Candy SG (2012) Channel properties reveal differential expression of TARPed and TARPless AMPARs in stargazer neurons. Nat Neurosci 15:853–861. doi: 10.1038/nn.3107.

Bellone C and Luscher C (2006) Cocaine triggered AMPA receptor redistribution is reversed in vivo by mGluR-dependent long-term depression. Nat Neurosci 9:636–641. doi: 10.1038/nn1682.

Benke T and Traynelis SF (2019) AMPA-Type Glutamate Receptor Conductance Changes and Plasticity: Still a Lot of Noise. Neurochem Res 44:539–548. doi: 10.1007/s11064-018-2491-1.

Bowie D (2012) Redefining the classification of AMPA-selective ionotropic glutamate receptors. J Physiol 590:49–61. doi: 10.1113/jphysiol.2011.221689.

Bowie D, Lange GD and Mayer ML (1998) Activity-dependent modulation of glutamate receptors by polyamines. J Neurosci 18:8175–8185. doi: 10.1523/jneurosci.18-20-08175.1998.

Bowie D and Mayer ML (1995) Inward rectification of both AMPA and kainate subtype glutamate receptors generated by polyamine-mediated ion channel block. Neuron 15:453–462. doi: 10.1016/0896-6273(95)90049-7.

Brill J and Huguenard JR (2008) Sequential changes in AMPA receptor targeting in the developing neocortical excitatory circuit. J Neurosci 28:13918–13928. doi: 10.1523/JNEUROSCI.3229-08.2008.

Brown P, McGuire H and Bowie D (2018) Stargazin and cornichon-3 relieve polyamine block of AMPA receptors by enhancing blocker permeation. J Gen Physiol 150:67–82. doi: 10.1085/jgp.201711895.

Brown PM, Aurousseau MR, Musgaard M, Biggin PC and Bowie D (2016) Kainate receptor pore-forming and auxiliary subunits regulate channel block by a novel mechanism. J Physiol 594:1821–1840. doi: 10.1113/JP271690.

Burnashev N, Monyer H, Seeburg PH and Sakmann B (1992) Divalent ion permeability of AMPA receptor channels is dominated by the edited form of a single subunit. Neuron 8:189–198. doi: 10.1016/0896-6273(92)90120-3.

Castilho A, Madsen E, Ambrosio AF, Veruki ML and Hartveit E (2015) Diabetic hyperglycemia reduces Ca2+ permeability of extrasynaptic AMPA receptors in AII amacrine cells. J Neurophysiol 114:1545–1553. doi: 10.1152/jn.00295.2015.

Cho CH, St-Gelais F, Zhang W, Tomita S and Howe JR (2007) Two families of TARP isoforms that have distinct effects on the kinetic properties of AMPA receptors and synaptic currents. Neuron 55:890–904. doi: 10.1016/j.neuron.2007.08.024.

Chung C, Deak F and Kavalali ET (2008) Molecular substrates mediating lanthanide-evoked neurotransmitter release in central synapses. J Neurophysiol 100:2089–2100. doi: 10.1152/jn.90404.2008.

Clem RL and Huganir RL (2010) Calcium-permeable AMPA receptor dynamics mediate fear memory erasure. Science 330:1108–1112. doi: 10.1126/science.1195298.

Conrad KL, Tseng KY, Uejima JL, Reimers JM, Heng LJ, Shaham Y, Marinelli M and Wolf ME (2008) Formation of accumbens GluR2-lacking AMPA receptors mediates incubation of cocaine craving. Nature 454:118–121. doi: 10.1038/nature06995.

Coombs ID, MacLean DM, Jayaraman V, Farrant M and Cull-Candy SG (2017) Dual Effects of TARP γ-2 on Glutamate Efficacy Can Account for AMPA Receptor Autoinactivation. Cell Rep 20:1123–1135. doi: 10.1016/j.celrep.2017.07.014.

Coombs ID, Soto D, Zonouzi M, Renzi M, Shelley C, Farrant M and Cull-Candy SG (2012) Cornichons modify channel properties of recombinant and glial AMPA receptors. J Neurosci 32:9796–9804. doi: 10.1523/JNEUROSCI.0345-12.2012.

Cull-Candy S, Kelly L and Farrant M (2006) Regulation of Ca2+-permeable AMPA receptors: synaptic plasticity and beyond. Curr Opin Neurobiol 16:288–297. doi: 10.1016/j.conb.2006.05.012.

Dixon RM, Mellor JR and Hanley JG (2009) PICK1-mediated glutamate receptor subunit 2 (GluR2) trafficking contributes to cell death in oxygen/glucose-deprived hippocampal neurons. J Biol Chem 284:14230–14235. doi: 10.1074/jbc.M901203200.

Donevan SD and Rogawski MA (1995) Intracellular polyamines mediate inward rectification of Ca(2+)-permeable alpha-amino-3-hydroxy-5-methyl-4-isoxazolepropionic acid receptors. Proc Natl Acad Sci U S A 92:9298–9302. doi: 10.1073/pnas.92.20.9298.

Fortin DA, Davare MA, Srivastava T, Brady JD, Nygaard S, Derkach VA and Soderling TR (2010) Long-term potentiation-dependent spine enlargement requires synaptic Ca2+-permeable AMPA receptors recruited by CaM-kinase I. J Neurosci 30:11565–11575. doi: 10.1523/JNEUROSCI.1746-10.2010.

Fukaya M, Yamazaki M, Sakimura K and Watanabe M (2005) Spatial diversity in gene expression for VDCCgamma subunit family in developing and adult mouse brains. Neurosci Res 53:376–383. doi: 10.1016/j.neures.2005.08.009.

Goffer Y, Xu D, Eberle SE, D’Amour J, Lee M, Tukey D, Froemke RC, Ziff EB and Wang J (2013) Calcium-permeable AMPA receptors in the nucleus accumbens regulate depression-like behaviors in the chronic neuropathic pain state. J Neurosci 33:19034–19044. doi: 10.1523/JNEUROSCI.2454-13.2013.

Gray EE, Fink AE, Sarinana J, Vissel B and O’Dell TJ (2007) Long-term potentiation in the hippocampal CA1 region does not require insertion and activation of GluR2-lacking AMPA receptors. J Neurophysiol 98:2488–2492. doi: 10.1152/jn.00473.2007.

Greger IH, Watson JF and Cull-Candy SG (2017) Structural and Functional Architecture of AMPA-Type Glutamate Receptors and Their Auxiliary Proteins. Neuron 94:713–730. doi: 10.1016/j.neuron.2017.04.009.

Huganir RL and Nicoll RA (2013) AMPARs and synaptic plasticity: the last 25 years. Neuron 80:704–717. doi: 10.1016/j.neuron.2013.10.025.

Jackson AC, Milstein AD, Soto D, Farrant M, Cull-Candy SG and Nicoll RA (2011) Probing TARP modulation of AMPA receptor conductance with polyamine toxins. J Neurosci 31:7511–7520. doi: 10.1523/JNEUROSCI.6688-10.2011.

Jackson AC and Nicoll RA (2011a) The expanding social network of ionotropic glutamate receptors: TARPs and other transmembrane auxiliary subunits. Neuron 70:178–199. doi: 10.1016/j.neuron.2011.04.007.

Jackson AC and Nicoll RA (2011b) Stargazin (TARP gamma-2) is required for compartment-specific AMPA receptor trafficking and synaptic plasticity in cerebellar stellate cells. J Neurosci 31:3939–3952. doi: 10.1523/JNEUROSCI.5134-10.2011.

Jia Z, Agopyan N, Miu P, Xiong Z, Henderson J, Gerlai R, Taverna FA, Velumian A, MacDonald J, Carlen P, Abramow-Newerly W and Roder J (1996) Enhanced LTP in mice deficient in the AMPA receptor GluR2. Neuron 17:945–956. doi: 10.1016/s0896-6273(00)80225-1.

Kamboj SK, Swanson GT and Cull-Candy SG (1995) Intracellular spermine confers rectification on rat calcium-permeable AMPA and kainate receptors. J Physiol 486 (Pt 2):297–303. doi: 10.1113/jphysiol.1995.sp020812.

Katano T, Furue H, Okuda-Ashitaka E, Tagaya M, Watanabe M, Yoshimura M and Ito S (2008) N-ethylmaleimide-sensitive fusion protein (NSF) is involved in central sensitization in the spinal cord through GluR2 subunit composition switch after inflammation. Eur J Neurosci 27:3161–3170. doi: 10.1111/j.1460-9568.2008.06293.x.

Koh DS, Burnashev N and Jonas P (1995) Block of native Ca(2+)-permeable AMPA receptors in rat brain by intracellular polyamines generates double rectification. J Physiol 486 (Pt 2):305–312. doi: 10.1113/jphysiol.1995.sp020813.

Koike M, Iino M and Ozawa S (1997) Blocking effect of 1-naphthyl acetyl spermine on Ca(2+)-permeable AMPA receptors in cultured rat hippocampal neurons. Neurosci Res 29:27–36. doi: 10.1016/s0168-0102(97)00067-9.

Korgaonkar AA, Li Y, Sekhar D, Subramanian D, Guevarra J, Swietek B, Pallottie A, Singh S, Kella K, Elkabes S and Santhakumar V (2020) Toll-like Receptor 4 Signaling in Neurons Enhances Calcium-Permeable alpha-Amino-3-Hydroxy-5-Methyl-4-Isoxazolepropionic Acid Receptor Currents and Drives Post-Traumatic Epileptogenesis. Ann Neurol 87:497–515. doi: 10.1002/ana.25698.

Kudoh SN and Taguchi T (2002) A simple exploratory algorithm for the accurate and fast detection of spontaneous synaptic events. Biosens Bioelectron 17:773–782. doi: 10.1016/s0956-5663(02)00053-2.

Kumar SS, Bacci A, Kharazia V and Huguenard JR (2002) A developmental switch of AMPA receptor subunits in neocortical pyramidal neurons. J Neurosci 22:3005–3015. doi: 10.1523/JNEUROSCI.22-08-03005.2002.

Kuner T, Beck C, Sakmann B and Seeburg PH (2001) Channel-lining residues of the AMPA receptor M2 segment: structural environment of the Q/R site and identification of the selectivity filter. J Neurosci 21:4162–4172. doi: 10.1523/JNEUROSCI.21-12-04162.2001.

Lamsa KP, Heeroma JH, Somogyi P, Rusakov DA and Kullmann DM (2007) Anti-Hebbian long-term potentiation in the hippocampal feedback inhibitory circuit. Science 315:1262–1266. doi: 10.1126/science.1137450.

Lee BR, Ma YY, Huang YH, Wang X, Otaka M, Ishikawa M, Neumann PA, Graziane NM, Brown TE, Suska A, Guo C, Lobo MK, Sesack SR, Wolf ME, Nestler EJ, Shaham Y, Schluter OM and Dong Y (2013) Maturation of silent synapses in amygdala-accumbens projection contributes to incubation of cocaine craving. Nat Neurosci 16:1644–1651. doi: 10.1038/nn.3533.

Liu S, Lau L, Wei J, Zhu D, Zou S, Sun HS, Fu Y, Liu F and Lu Y (2004) Expression of Ca(2+)-permeable AMPA receptor channels primes cell death in transient forebrain ischemia. Neuron 43:43–55. doi: 10.1016/j.neuron.2004.06.017.

Liu SJ and Cull-Candy SG (2002) Activity-dependent change in AMPA receptor properties in cerebellar stellate cells. J Neurosci 22:3881–3889. doi: 10.1523/JNEUROSCI.22-10-03881.2002.

Liu SQ and Cull-Candy SG (2000) Synaptic activity at calcium-permeable AMPA receptors induces a switch in receptor subtype. Nature 405:454–458. doi: 10.1038/35013064.

Liu Y, Formisano L, Savtchouk I, Takayasu Y, Szabo G, Zukin RS and Liu SJ (2010) A single fear-inducing stimulus induces a transcription-dependent switch in synaptic AMPAR phenotype. Nat Neurosci 13:223–231. doi: 10.1038/nn.2474.

Lu W, Shi Y, Jackson AC, Bjorgan K, During MJ, Sprengel R, Seeburg PH and Nicoll RA (2009) Subunit composition of synaptic AMPA receptors revealed by a single-cell genetic approach. Neuron 62:254–268. doi: 10.1016/j.neuron.2009.02.027.

Lujan B, Dagostin A and von Gersdorff H (2019) Presynaptic Diversity Revealed by Ca(2+)-Permeable AMPA Receptors at the Calyx of Held Synapse. J Neurosci 39:2981–2994. doi: 10.1523/JNEUROSCI.2565-18.2019.

Magazanik LG, Buldakova SL, Samoilova MV, Gmiro VE, Mellor IR and Usherwood PN (1997) Block of open channels of recombinant AMPA receptors and native AMPA/kainate receptors by adamantane derivatives. J Physiol 505 (Pt 3):655–663. doi: 10.1111/j.1469-7793.1997.655ba.x.

Mahanty NK and Sah P (1998) Calcium-permeable AMPA receptors mediate long-term potentiation in interneurons in the amygdala. Nature 394:683–687. doi: 10.1038/29312.

Mainen ZF, Jia Z, Roder J and Malinow R (1998) Use-dependent AMPA receptor block in mice lacking GluR2 suggests postsynaptic site for LTP expression. Nat Neurosci 1:579–586. doi: 10.1038/2812.

McGee TP, Bats C, Farrant M and Cull-Candy SG (2015) Auxiliary Subunit GSG1L Acts to Suppress Calcium-Permeable AMPA Receptor Function. J Neurosci 35:16171–16179. doi: 10.1523/JNEUROSCI.2152-15.2015.

Nilsen A and England PM (2007) A subtype-selective, use-dependent inhibitor of native AMPA receptors. J Am Chem Soc 129:4902–4903. doi: 10.1021/ja0705801.

Ozawa S, Iino M and Tsuzuki K (1991) Two types of kainate response in cultured rat hippocampal neurons. J Neurophysiol 66:2–11. doi: 10.1152/jn.1991.66.1.2.

Panchenko VA, Glasser CR, Partin KM and Mayer ML (1999) Amino acid substitutions in the pore of rat glutamate receptors at sites influencing block by polyamines. J Physiol 520 Pt 2:337–357. doi: 10.1111/j.1469-7793.1999.t01-1-00337.x.

Park JS, Voitenko N, Petralia RS, Guan X, Xu JT, Steinberg JP, Takamiya K, Sotnik A, Kopach O, Huganir RL and Tao YX (2009) Persistent inflammation induces GluR2 internalization via NMDA receptor-triggered PKC activation in dorsal horn neurons. J Neurosci 29:3206–3219. doi: 10.1523/JNEUROSCI.4514-08.2009.

Peng PL, Zhong X, Tu W, Soundarapandian MM, Molner P, Zhu D, Lau L, Liu S, Liu F and Lu Y (2006) ADAR2-dependent RNA editing of AMPA receptor subunit GluR2 determines vulnerability of neurons in forebrain ischemia. Neuron 49:719–733. doi: 10.1016/j.neuron.2006.01.025.

Plant K, Pelkey KA, Bortolotto ZA, Morita D, Terashima A, McBain CJ, Collingridge GL and Isaac JT (2006) Transient incorporation of native GluR2-lacking AMPA receptors during hippocampal long-term potentiation. Nat Neurosci 9:602–604. doi: 10.1038/nn1678.

Poulsen MH, Lucas S, Stromgaard K and Kristensen AS (2014) Evaluation of PhTX-74 as subtype-selective inhibitor of GluA2-containing AMPA receptors. Mol Pharmacol 85:261–268. doi: 10.1124/mol.113.089961.

Rothman JS and Silver RA (2018) NeuroMatic: An Integrated Open-Source Software Toolkit for Acquisition, Analysis and Simulation of Electrophysiological Data. Front Neuroinform 12:14. doi: 10.3389/fninf.2018.00014.

Rozov A, Zilberter Y, Wollmuth LP and Burnashev N (1998) Facilitation of currents through rat Ca2+-permeable AMPA receptor channels by activity-dependent relief from polyamine block. J Physiol 511 (Pt 2):361–377. doi: 10.1111/j.1469-7793.1998.361bh.x.

Sanderson JL, Gorski JA and Dell’Acqua ML (2016) NMDA Receptor-Dependent LTD Requires Transient Synaptic Incorporation of Ca(2)(+)-Permeable AMPARs Mediated by AKAP150-Anchored PKA and Calcineurin. Neuron 89:1000–1015. doi: 10.1016/j.neuron.2016.01.043.

Sara Y, Bal M, Adachi M, Monteggia LM and Kavalali ET (2011) Use-dependent AMPA receptor block reveals segregation of spontaneous and evoked glutamatergic neurotransmission. J Neurosci 31:5378–5382. doi: 10.1523/JNEUROSCI.5234-10.2011.

Scheyer AF, Wolf ME and Tseng KY (2014) A protein synthesis-dependent mechanism sustains calcium-permeable AMPA receptor transmission in nucleus accumbens synapses during withdrawal from cocaine self-administration. J Neurosci 34:3095–3100. doi: 10.1523/JNEUROSCI.4940-13.2014.

Schwenk J, Boudkkazi S, Kocylowski MK, Brechet A, Zolles G, Bus T, Costa K, Kollewe A, Jordan J, Bank J, Bildl W, Sprengel R, Kulik A, Roeper J, Schulte U and Fakler B (2019) An ER Assembly Line of AMPA-Receptors Controls Excitatory Neurotransmission and Its Plasticity. Neuron 104:680–692 e689. doi: 10.1016/j.neuron.2019.08.033.

Sladek AL and Nawy S (2020) Ocular Hypertension Drives Remodeling of AMPA Receptors in Select Populations of Retinal Ganglion Cells. Front Synaptic Neurosci 12:30. doi: 10.3389/fnsyn.2020.00030.

Sommer B, Kohler M, Sprengel R and Seeburg PH (1991) RNA editing in brain controls a determinant of ion flow in glutamate-gated channels. Cell 67:11–19. doi: 10.1016/0092-8674(91)90568-j.

Soto D, Coombs ID, Gratacos-Batlle E, Farrant M and Cull-Candy SG (2014) Molecular mechanisms contributing to TARP regulation of channel conductance and polyamine block of calcium-permeable AMPA receptors. J Neurosci 34:11673–11683. doi: 10.1523/JNEUROSCI.0383-14.2014.

Soto D, Coombs ID, Kelly L, Farrant M and Cull-Candy SG (2007) Stargazin attenuates intracellular polyamine block of calcium-permeable AMPA receptors. Nat Neurosci 10:1260–1267. doi: 10.1038/nn1966.

Soto D, Coombs ID, Renzi M, Zonouzi M, Farrant M and Cull-Candy SG (2009) Selective regulation of long-form calcium-permeable AMPA receptors by an atypical TARP, gamma-5. Nat Neurosci 12:277–285. doi: 10.1038/nn.2266.

Studniarczyk D, Coombs I, Cull-Candy SG and Farrant M (2013) TARP gamma-7 selectively enhances synaptic expression of calcium-permeable AMPARs. Nat Neurosci 16:1266–1274. doi: 10.1038/nn.3473.

Sullivan SJ, Farrant M and Cull-Candy SG (2017) TARP gamma-2 Is Required for Inflammation-Associated AMPA Receptor Plasticity within Lamina II of the Spinal Cord Dorsal Horn. J Neurosci 37:6007–6020. doi: 10.1523/JNEUROSCI.0772-16.2017.

Swanson GT, Kamboj SK and Cull-Candy SG (1997) Single-channel properties of recombinant AMPA receptors depend on RNA editing, splice variation, and subunit composition. J Neurosci 17:58–69. doi: 10.1523/JNEUROSCI.17-01-00058.1997.

Tikhonov DB (2007) Ion channels of glutamate receptors: structural modeling. Mol Membr Biol 24:135–147. doi: 10.1080/09687860601008806.

Tikhonov DB, Samoilova MV, Buldakova SL, Gmiro VE and Magazanik LG (2000) Voltage-dependent block of native AMPA receptor channels by dicationic compounds. Br J Pharmacol 129:265–274. doi: 10.1038/sj.bjp.0703043.

Toth K and McBain CJ (1998) Afferent-specific innervation of two distinct AMPA receptor subtypes on single hippocampal interneurons. Nat Neurosci 1:572–578. doi: 10.1038/2807.

Traynelis SF, Wollmuth LP, McBain CJ, Menniti FS, Vance KM, Ogden KK, Hansen KB, Yuan H, Myers SJ and Dingledine R (2010) Glutamate receptor ion channels: structure, regulation, and function. Pharmacol Rev 62:405–496. doi: 10.1124/pr.109.002451.

Tsubokawa H, Oguro K, Masuzawa T, Nakaima T and Kawai N (1995) Effects of a spider toxin and its analogue on glutamate-activated currents in the hippocampal CA1 neuron after ischemia. J Neurophysiol 74:218–225. doi: 10.1152/jn.1995.74.1.218.

Twomey EC, Yelshanskaya MV, Vassilevski AA and Sobolevsky AI (2018) Mechanisms of Channel Block in Calcium-Permeable AMPA Receptors. Neuron 99:956–968 e954. doi: 10.1016/j.neuron.2018.07.027.

Van den Oever MC, Goriounova NA, Li KW, Van der Schors RC, Binnekade R, Schoffelmeer AN, Mansvelder HD, Smit AB, Spijker S and De Vries TJ (2008) Prefrontal cortex AMPA receptor plasticity is crucial for cue-induced relapse to heroin-seeking. Nat Neurosci 11:1053–1058. doi: 10.1038/nn.2165.

Vikman KS, Rycroft BK and Christie MJ (2008) Switch to Ca2+-permeable AMPA and reduced NR2B NMDA receptor-mediated neurotransmission at dorsal horn nociceptive synapses during inflammatory pain in the rat. J Physiol 586:515–527. doi: 10.1113/jphysiol.2007.145581.

Wang HX and Gao WJ (2010) Development of calcium-permeable AMPA receptors and their correlation with NMDA receptors in fast-spiking interneurons of rat prefrontal cortex. J Physiol 588:2823–2838. doi: 10.1113/jphysiol.2010.187591.

Washburn MS and Dingledine R (1996) Block of alpha-amino-3-hydroxy-5-methyl-4-isoxazolepropionic acid (AMPA) receptors by polyamines and polyamine toxins. J Pharmacol Exp Ther 278:669–678.

Weiss NA (2016) wBoot: Bootstrap Methods. R package version 1.0.3. https://CRAN.R-project.org/package=wBoot.

Wenthold RJ, Petralia RS, Blahos J, II and Niedzielski AS (1996) Evidence for multiple AMPA receptor complexes in hippocampal CA1/CA2 neurons. J Neurosci 16:1982–1989. doi: 10.1523/JNEUROSCI.16-06-01982.1996.

Yamasaki M, Miyazaki T, Azechi H, Abe M, Natsume R, Hagiwara T, Aiba A, Mishina M, Sakimura K and Watanabe M (2011) Glutamate receptor delta2 is essential for input pathway-dependent regulation of synaptic AMPAR contents in cerebellar Purkinje cells. J Neurosci 31:3362–3374. doi: 10.1523/JNEUROSCI.5601-10.2011.

Yamazaki M, Fukaya M, Hashimoto K, Yamasaki M, Tsujita M, Itakura M, Abe M, Natsume R, Takahashi M, Kano M, Sakimura K and Watanabe M (2010) TARPs gamma-2 and gamma-7 are essential for AMPA receptor expression in the cerebellum. Eur J Neurosci 31:2204–2220. doi: 10.1111/j.1460-9568.2010.07254.x.

Zaitsev AV, Kim KK, Fedorova IM, Dorofeeva NA, Magazanik LG and Tikhonov DB (2011) Specific mechanism of use-dependent channel block of calcium-permeable AMPA receptors provides activity-dependent inhibition of glutamatergic neurotransmission. J Physiol 589:1587–1601. doi: 10.1113/jphysiol.2011.204362.

Zhao Y, Chen S, Swensen AC, Qian WJ and Gouaux E (2019) Architecture and subunit arrangement of native AMPA receptors elucidated by cryo-EM. Science 364:355–362. doi: 10.1126/science.aaw8250.

